# Compensation of physiological motion enables high-yield whole-cell recording *in vivo*

**DOI:** 10.1101/2020.06.09.143008

**Authors:** William M. Stoy, Bo Yang, Ali Kight, Nathaniel C. Wright, Peter Y. Borden, Garrett B. Stanley, Craig R. Forest

## Abstract

Whole-cell patch-clamp recording *in vivo* is the gold-standard method for measuring subthreshold electrophysiology from single cells during behavioural tasks, sensory stimulations, and optogenetic manipulation. However, these recordings require a tight, gigaohm resistance, seal between a glass pipette electrode’s aperture and a cell’s membrane. These seals are difficult to form, especially in vivo, in part because of a strong dependence on the distance between the pipette aperture and cell membrane. We elucidate and utilize this dependency to develop an autonomous method for placement and synchronization of pipette’s tip aperture to the membrane of a nearby, moving neuron, which enables high-yield seal formation and subsequent recordings in the deep in the brain of the living mouse, in the thalamus. This synchronization procedure nearly doubles the reported gigaseal yield in the thalamus (>3 mm below the pial surface) from 26% (n=17/64) to 48% (n=32/66). Whole-cell recording yield improved from 10% (n = 9/88) to 24% (n=18/76) when motion compensation was used during the gigaseal formation. As an example of its application, we utilized this system to investigate the role of the sensory environment and ventral posterior medial region (VPM) projection synchrony on intracellular dynamics in the barrel cortex. This method results in substantially greater subcortical whole-cell recording yield than previously reported and thus makes pan-brain whole-cell electrophysiology practical in the living mouse brain.

## 1.1.2 Introduction

Physiological motion due to, for example, breathing and heart beating, presents a significant barrier to reliable measurement of *in vivo* neural phenomena. Intracellular probes are especially sensitive to this displacement, since a cell’s diameter and motion amplitude are comparable. For example, although sharp intracellular recordings were developed in 1949 by Gerard and Ling, recordings longer than several seconds have been historically difficult to obtain, due primarily to micromotion from the heartbeat and respiration of the animal (Britt & Rossi, 1982b). This resulted in only a handful of papers describing the use of sharp intracellular recordings *in vivo* in over half a century since its development (Bazhenov et al., 1999; Kim & McCormick, 1998; Lampl et al., 1999; Ylinen et al., 1995), relegating the method primarily to *in vitro* use. While whole-cell patch clamping solves this problem with the robust mechanical stability of the gigaohm seal between the pipette and the cell’s membrane, it has been previously noted that micromotion can still present a challenge for the initial formation of this stable configuration (Margrie et al., 2002).

The reported amplitudes of this physiological micromotion in the brains of rodents are varied. Lee et al. report approximately 1 µm peak-to-peak for cardiac induced motion, and 10 µm for respiratory induced motion in anesthetized mice (Lee, Courties, et al., 2017). Gilletti and Muthuswamy report 5-10 µm average for breathing motion and 2-3 µm average for heartbeat in anesthetized Sprague Dawley rats (Gilletti & Muthuswamy, 2006). Fee reports an approximately 2 and 4 µm peak to peak average for cardiac and respiratory motion, respectively, in awake rats (Fee, 2000).

A variety of strategies have arisen to measure this motion in real-time, including laser interferometry and laser triangulation (Fee, 2000; Lee, Vinegoni, et al., 2017), as well as high-speed motion detection and active stabilization of the objective lens (Lee, Nakamura, et al., 2008). However, these techniques are limited to superficial layers due to the highly scattering optical properties of brain tissue. Others have developed motion transducers for placement on the brain surface (Gilletti & Muthuswamy, 2006), and transducer probes inserted into the brain (Kursu et al., 2012; Lee, Ozaki, et al., 2008; Vähäsöyrinki et al., 2009). However, these methods are not applicable for patch clamp where only the micromotion of the tissue near the tip of the pipette is relevant. Still others have suggested invasive surgical procedure to reduce respiration and cardiac motion artifacts *in vivo*, such as draining the spinal fluid or opening the chest cavity (Britt & Rossi, 1982a, 1982b; Matsumoto et al., 2011). An ideal solution would preserve the physiological relevance of the preparation and be applicable to whole-cell patch recording throughout the brain, from the cortical surface to deep, subcortical nuclei.

Accordingly, we have developed a system to measure the local micromotion of cells with respect to the patch pipette tip that uses the series resistance to ground as the measurement of cell-membrane distance. With this system, no external measurements of brain motion need be utilized, and no invasive surgical procedures are required to measure the heartbeat or breathing. Additionally, we demonstrate a feedforward position control system that can compensate for this micromotion, by reducing the amplitude of the fluctuations of this electrical resistance, and thus improve the success rate of gigaseal formation and subsequent whole-cell patch clamp recording.

We first characterize the sensitivity of the gigaseal formation to the pipette-membrane distance by attempting a gigaseal with the patch pipette tip placed at various distances with respect to the surface of plated HEK293 cells *in vitro*. We found that there is a very small window near the surface of the cell where gigaseal has a high probability of success, approximately 1-3 µm.

We then recorded and compensated for micromotion in the brain. The motion due to the heartbeat and breathing moves cells in the brain with respect to the pipette tip. When a cell’s membrane is near the tip of the pipette, it blocks the flow of current to the ground electrode in the saline bath above the animal’s head. Therefore, the oscillatory modulation of cell’s position is detectable as correlated oscillatory modulation (fluctuations) of the electrode current measured in voltage clamp mode on the patch amplifier.

We designed a system to use this changing impedance as a measure of the distance to the cell’s membrane and predict the motion caused by the animal’s heartbeat and breathing by computing the finite impulse response from each cardiac and respiratory event, conceptually related to the work of Fee in anesthetized zebra finches with sharp intracellular electrodes (Fee, 2000). The resulting system is capable of reducing the relative motion of the patch pipette with respect to a nearby cell, which, when suction is applied, improves gigaseal and whole-cell yield in the cortex, and approximately doubles the probability of achieving a gigaseal and whole-cell state in the thalamus.

## 1.1 Methods

### 1.1.3 Acute in vivo preparation

All experiments were performed in accordance with the Georgia Tech Institutional Animal Care and Use Committee (IACUC) guidelines. For *in vivo* preparation, all mice (n=17) were prepared for acute experimentation (Stoy et al., 2017). Briefly, male C57BL/6 mice (p35–p49) were anesthetized with isoflurane and headfixed to a titanium headplate with C&B-Metabond dental cement (Parkell, Edgewood, NY). Craniotomies (1 mm diameter) and durotomies were performed above the Ventral Posteromedial nucleus (VPM) of the thalamus (1.75 mm Rostral, 1.75 mm Lateral, 2.7-3.3 mm below the pial surface). For cortical experiments, craniotomies (0.5 mm in diameter) were performed above the Barrel Cortex (1.8 mm rostral, 2.6 mm lateral, 400 µm below the pial surface). In both preparations, stereotaxic coordinates from the Paxinos and Franklin mouse brain atlas (Paxinos & Franklin, 2012) were used. Patch-clamp experiments were performed in aCSF consisting of (in mM): 135 NaCl, 5 HEPES, 1 MgCl2, 5 KCl, 1.8 CaCl2 (pH: 7.3, osmolarity: approximately 285 mOsm). The internal pipette solution consisted of (in mM): 135 K Gluconate, 10 HEPES, 4 KCl, 1 EGTA, 0.3 GTP-Na, 4 ATP-Mg, 10 Na2-Phosphocreatine (pH 7.25, osmolarity: approximately 290 mOsm). All pipettes were pulled from standard wall, filamented, borosilicate glass (GC150F-10, Warner Instruments) to a resistance between 6 and 10 MΩ.

### 1.1.4 HEK293T cell preparation

Human embryonic kidney (HEK293T) cells (American Type Culture Collection, Manassas, VA) were maintained at 37°C and 5% CO_2_ in Eagle Minimum Essential Medium (MEM) supplemented with 5% FBS, 40μM L-glutamine, 100 U/ml penicillin and 0.1 mM streptomycin. Cells were passaged regularly and split when they reached 70% confluency. Cells used for *in vitro* electrophysiology experiments were grown on glass coverslips (12×12mm, No.2, VWR). Patch-clamp experiments were performed in aCSF consisting of (in mM): 161 NaCl, 10 HEPES, 6 D-Glucose, 3 KCl, 1 MgCl2, 1.5 CaCl2 (pH: 7.4). The internal pipette solution consisted of (in mM): 120 KCl, 2 MgCl2, 10 EGTA, 10 HEPES (pH: 7.2-7.3, osmolarity: 290-300 mOsm).

### 1.1.5 Hardware and setup

The hardware setup is shown in Figure 1 and consists of several sub-systems, including the physiology, impulse response calculation, and patch clamp subsystems. The patch clamp subsystem itself consists of manipulator, electrophysiology, and pressure control subsystems.

**Figure 1:**
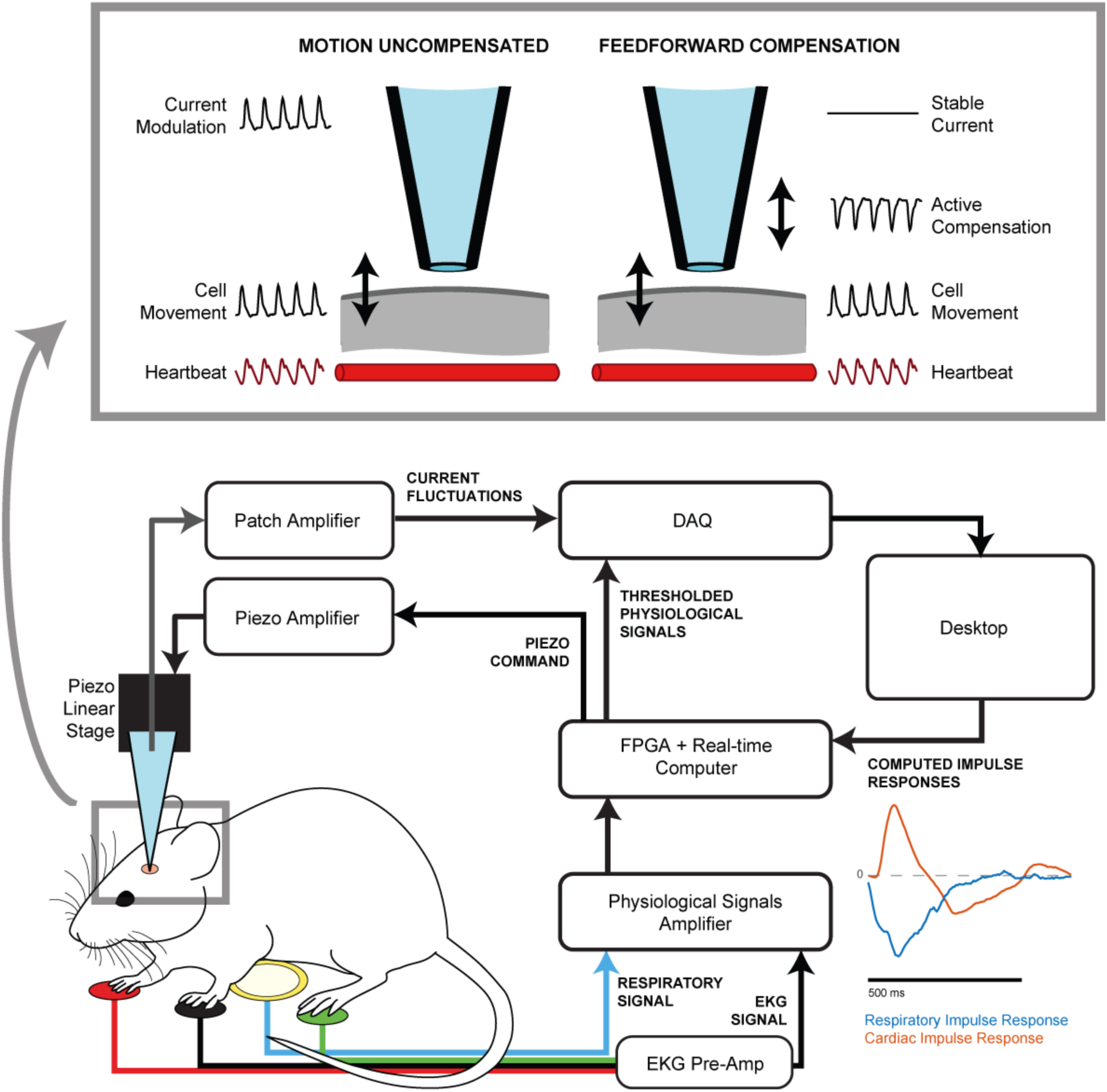
Schematic of physiological motion compensation instrumentation and method for high-yield whole-cell patch clamp electrophysiology *in vivo*. Cardiac signals are recorded with an electrocardiogram (EKG) amplifier and respiratory signals are captured with a piezo pickup under the animal’s chest. These physiological signals are amplified and thresholded on a Field Programmable Gate Array (FPGA). Simultaneously, current fluctuations, indicative of a nearby cell moving with respect to the pipette tip are amplified on a patch amplifier and recorded along with physiological signals on a data acquisition device (DAQ). A desktop computer then computes the cardiac and respiratory impulse responses and sends them to the real-time computer. A real-time computer computes the appropriate command signal required to synchronize the position of the pipette tip with respect to the cell by scaling and convolving the impulse responses with the thresholded physiological signals. The command signal is then amplified by a piezo amplifier and is used to drive a piezo stack in a piezo linear stage. The relative motion between the pipette and the cell is compensated by the motion of the piezo stage, resulting in the measurement of a stable current, and hence no relative motion, between the pipette electrode and cell membrane in front of it.

### 1.1.6 Cardiac and respiratory physiological recordings

The heartbeat was recorded using an EKG preamplifier (C-ISO-256) and a Two Channel Combination Bridge EKG amplifier (ETH-256, iworx). Leads were connected to the animal’s front paws and right hind paw via Skintact FS-TB1-5-Gel ECG Electrode (VWR) as shown in Figure 1.

The cardiac signal was amplified 10x and a 0.03 Hz – 10 kHz band pass filter was applied. Respiration was recorded non-invasively by placing a 20mm guitar pickup piezo disc element underneath the mouse’s chest. The respiratory signal was scaled by 50x and a 0.3 Hz – 10 kHz band pass filter was applied. Cardiac and respiratory signals were thresholded using a Schmitt trigger configuration with a time lockout to prevent triggering on noise in the input signals, implemented on the myRIO FPGA. The threshold level was set manually at the beginning of the experiment by adjusting a slider in the GUI. The thresholded physiological signals and the primary output (patch headstage in voltage clamp mode) from the Multiclamp 700B were recorded for impulse response estimation on an NI USB-6221 BNC DAQ (National Instruments) and also recorded on a Molecular Devices Digidata 1550 digitizer and visualized with PClamp 10 (Molecular Devices, Sunnyvale, CA) for post processing.

### 1.1.7 Patch clamp system

#### 1.1.7.1 Manipulator subsystem

Gross manipulation of the pipette is performed under manual control of the Sutter MP 285 3-axis patch manipulators (Sutter Instruments). Axial movements of the pipette are performed by the one-axis motorized translation stage (PT1-Z8, Thorlabs) is mounted to the z-stage of the Sutter Manipulator at a 45° angle from the vertical. A single axis linear piezo flexure stage (P-611.1, Phisik Instrumente) with 100 µm travel range mounted parallel to the Thorlabs stage for axial motion synchronization of the pipette tip with respect to the cell membrane. The pipette holder was attached to the piezo stage using a custom machined aluminum bracket.

#### 1.1.7.2 *In vivo* electrophysiology and pressure control subsystems

Whole-cell patch clamping was performed as described previously (Kodandaramaiah et al., 2016). Briefly, an Autopatcher 1500 (Neuromatic Devices) was used to provide computer-controlled pressure and measure resistance for both *in vitro* and *in vivo* experiments. Both *in vitro* and *in vivo* experiments were performed with Multiclamp 700B amplifiers (Molecular Devices) and signals were digitized at 20kHz (cDAQ-9174, National Instruments), and recorded in PClamp 10. All whole-cell recordings were performed in current clamp mode.

As in our previous work (Stoy et al., 2017) where we demonstrated an algorithm for navigation around blood vessels during regional pipette localization (RPL), resistance measurements were performed throughout RPL, rather than only before and after localization. Briefly, during localization to the region of interest (cortex or thalamus for this study), the pipette resistance was recorded in the aCSF on the surface of the tissue by applying a 20 mV amplitude, 128 Hz square wave at 50% duty cycle and stored as a moving average of four periods. If an obstruction was encountered during RPL (noted by an increase in resistance above a threshold), a sequence of motion and pressure control steps were issued to navigate electrodes efficiently around putative blood vessels, return to the original pipette axis, and continue neuron hunting.

### 1.1.8 Software

#### 1.1.8.1 Synchronization procedure

Neuron hunting and neuron detection proceeds as discussed in our previous work (Stoy et al., 2017). When a neuron is encountered during the neuron hunting step, forward motion of the pipette (here, 1 µm/s) is halted, the differential resistance square wave is stopped, and the synchronization procedure begins.

The relationship between a dynamic system’s inputs (here, the distance between the pipette and the cell), and its inputs (here, the cardiac and respiratory events) is defined as the system’s impulse response. The dynamic system comprises the mechanical transmission of movement from the heartbeats and breaths to the movement of the cell and is assumed to be linear and time invariant (LTI). As such, the system is completely characterized by the impulse response, and a prediction of the output can be made by convolving the system’s input signals with its impulse response. This prediction is used to drive the position of the pipette in concert with the motion of the cell.

A proxy measurement for the pipette-cell distance is made by applying a -65 mV holding voltage to the pipette in voltage clamp mode, enabling high temporal resolution recording of the pipette current, which is inversely proportional to the DC pipette resistance. The current from the amplifier and the thresholded cardiac and respiratory signals are recorded simultaneously in a ring buffer for 8 seconds. The signals were decimated to 200 Hz to improve the speed of computation. The impulse response was calculated using the MATLAB function *impulseest*. This function can take one or several input arrays and one output array as arguments and performs a non-parametric impulse response estimation of a given order using a recursive least squares estimation. The resulting cardiac and respiratory impulse responses were delivered to the myRIO embedded real-time system via NI shared variable synchronization over USB. The impulse responses were then interpolated to 1 kHz and convolved with the rising edges of the thresholded heartbeat and respiration signals on the real-time system using custom LabVIEW code on the myRIO real-time system, resulting in an estimation of the motion of the cell with respect to the pipette. This motion estimation was used as a command signal to drive the position of the piezo stage. The command signal was scaled manually by monitoring the fluctuations on the current trace until a suitable reduction in amplitude was observed, indicating sufficient synchronization of the position of the pipette and cell to perform a gigaseal attempt. The gigaseal attempt is queued by user input and is performed following the measurement of the next breath. When the next rising edge from the thresholded respiratory signal is detected, a gigasealing and break-in procedure are attempted after a 0.5 second delay as described below. During the gigasealing procedure, if the pipette resistance (Rp) crosses a 300 MΩ threshold, the amplitude of the piezo stage command signal is reduced linearly over 3 sec to 0 V. If the gigasealing procedure is aborted, the piezo command signal is immediately reduced to 0 V.

#### 1.1.8.2 Gigasealing and break-in procedure

When the gigasealing procedure begins, the system restarts the differential measurement of pipette resistance by applying a 20 mV square voltage pulse, and -40 mbar of suction is applied to the pipette. After 15 s of suction, the pressure is switched to atmospheric and the system will wait for the establishment of a gigaseal, defined as a measurement of total resistance over 1000 MΩ. Following formation of the gigaseal, the differential resistance square wave is stopped, and the fast and slow pipette and patch capacitance are compensated.

The break-in procedure is performed by applying a series of up to 15 pulses of suction at -345 mbar for increasing durations. Because the suction pressure is capacitively filtered by the tubing leading from the regulator to the pipette holder, increasing the suction time has the effect of also increasing the maximum pressure that is applied to the patch of membrane in the pipette. The break-in duration is increased by 25 ms prior to each attempt. Following each attempt, the program waits 1.25 s and averages the last 10 resistance measurements. If the resistance is above 800 MΩ, the program will immediately perform another break-in attempt as described above. If the resistance is between 300 MΩ and 800 MΩ, the program will wait until the seal has ‘recovered’ to 800 MΩ and then perform another break-in attempt. If the resistance is below 300 MΩ, the program assumes that a break-in has occurred and returns control to the user for recording. At this point, the user may choose to perform maintenance on the patch (apply slight negative suction to counteract the pressure head from the fluid in the pipette, or attempt to further break into the cell manually, etc.) or the user may proceed onto electrophysiological recording of the whole-cell. After 60 s, or manual abort of the break-in process, the pipette is retracted to the surface of the brain to be cleaned as we have described before (I Kolb et al., 2016) or exchanged if desired.

### 1.1.9 Measurement of in vitro relationship between gigaseal yield and pipette-membrane distance

The sensitivity of gigaseal formation to the distance between the membrane and the pipette tip was determined *in vitro* using HEK293T cells. The cells were visualized under 40X magnification using DIC optics. For each trial, a pipette filled with intracellular solution under 30 mbar of positive pressure was lowered near the surface of a cell and centered in X and Y above it. The pipette was then stepped down in Z in 200 nm steps and the resistance was measured after each step. When the pipette resistance had increased 0.1 MΩ above the initial pipette resistance (the threshold to begin gigasealing as defined in our previous work (I Kolb et al., 2016)), the Z position of the manipulator was defined as the “cell surface”. The downward pipette stepping was halted when the resistance had increased to 0.2 MΩ above the initial pipette resistance. The pipette was then retracted 30 µm (to eliminate backlash) and then moved to a random Z position between 3 µm above the cell surface to 7 µm below the cell surface. The positive pressure was then immediately released and a gigaseal and break-in procedure is performed as described previously (I Kolb et al., 2016). The success or failure of the gigaseal and subsequent whole-cell state was recorded.

### 1.1.10 Application of high yield autopatching techniques to an investigation of the thalamocortical pathway

In a subset of cortical experiments (n=2 mice), we applied the motion compensation technique described in this work, as well as robotic navigation during RPL (Stoy et al., 2017), and pipette cleaning (I Kolb et al., 2016; Ilya Kolb et al., 2019), towards an investigation of the mouse thalamocortical pathway. We utilized this system to investigate the role of the sensory environment and ventral posterior medial region (VPM) projection synchrony on intracellular dynamics in the barrel cortex.

A transgenic mouse colony was developed that expresses the opsin ChR2 in the VPM (See Supplementary Section S1.2). These neurons send excitatory projections to the barrel cortex (layer 4), which are stimulated using a fiber-coupled blue LED (470 nm, Thorlabs). This optical stimulation was combined with whisker stimulation from a galvanometer. The barrel cortex of each mouse was mapped using intrinsic signal optical imaging (ISOI) to identify regions representing the C1 and C2 whisker (Borden et al., 2017).

## 1.2 Results

### 1.1.11 Effect of pipette-membrane distance on gigasealing yield in vitro

To explore the relationship between small changes in the distance between the pipette and cell membrane and the probability of gigaseal formation, we performed an experiment in HEK293T cells. On each trial, the distance between the pipette and the membrane was randomly selected to be between 3 µm above and 7 µm below the surface of the cell. At each location, positive pressure on the patch pipette was released and the formation of a gigaseal was attempted on the cell (n=59 attempts, 40 gigaseals, Figure 2a). The resulting yield (successful gigaseals / number of attempts) with respect to pipette-membrane distance was smoothed with a 2 µm moving average window at 0.1 µm intervals. At the surface of the cell, 0 µm on Figure 2b, indicated by a vertical line, the gigaseal yield is 75% (n=9 gigaseals / 12 attempts).

**Figure 2:**
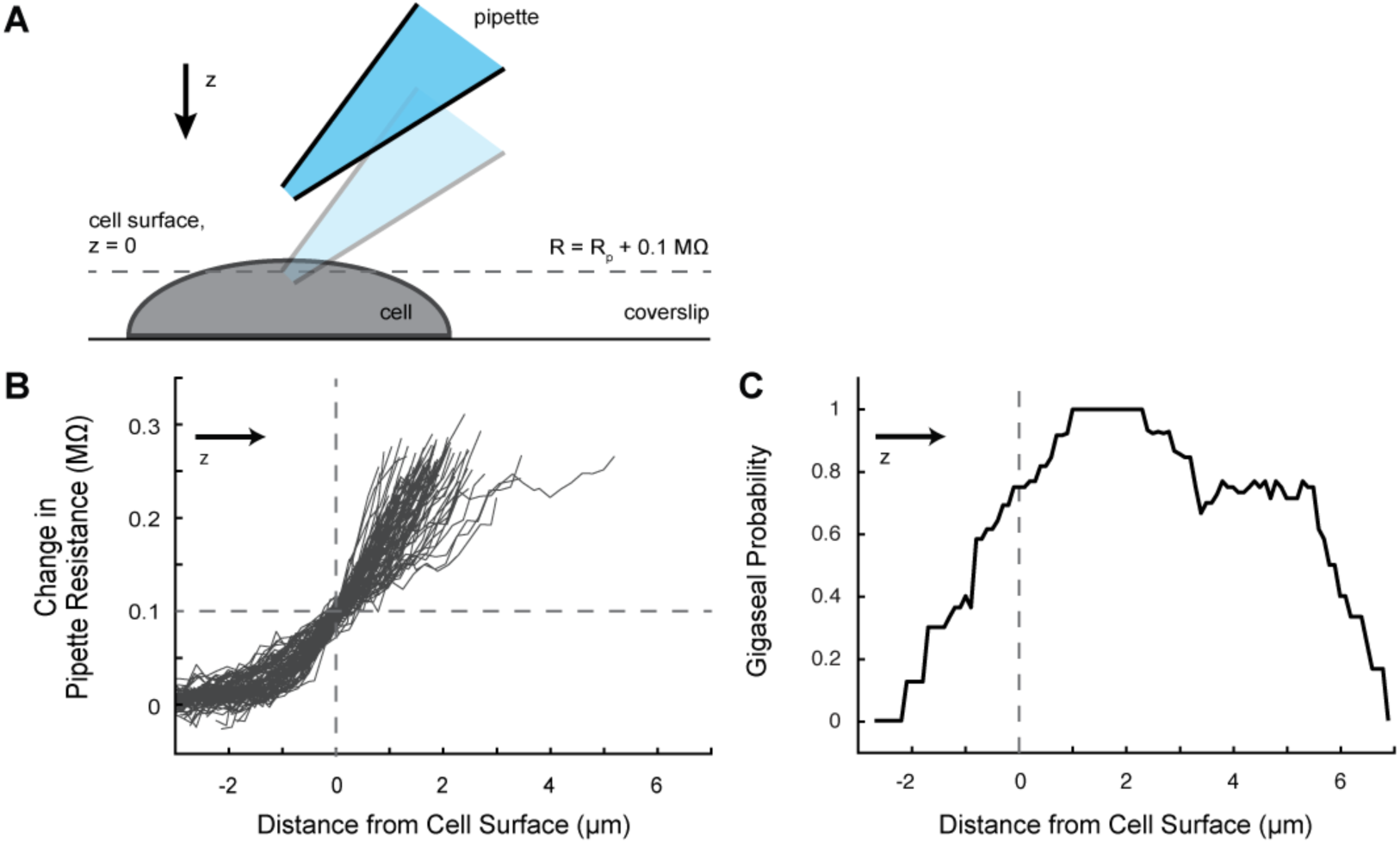
Measurement of the relationship between pipette-membrane distance and gigaseal probability. A pipette was slowly lowered vertically onto a plated HEK293T cell. At a resistance increase of 0.1 MΩ, the pipette is determined to be at the surface of the cell. The pipette is then moved into a random position between 3 µm above and 7 µm below the surface of the cell and a gigaseal is attempted. A) shows a schematic of this process. B) Pipette resistance increases as the pipette is stepped down into the cell. The cell surface is defined as the location where the pipette resistance increases 0.1 MΩ above its initial resistance in the saline bath far above the cell. C) Probability of gigaseal formation as a function of pipette position with respect to the membrane. Yield was calculated as the number of gigaseals / total number of attempts within a sliding 2 µm window. Pipettes 2.5 µm or more above the cell surface did not form a gigaseal with the pipette. Yield was maximized 1-2 µm below the surface of the cell.

The results in this experiment results conform to our previous measurements of the probability of achieving a whole-cell configuration in HEK293 cells (p=1, n=54/70, Fisher’s Exact Test, (Ilya Kolb et al., 2019)), although break-in was not attempted on the cells in this experiment. Within a 3 µm window (2 µm above the cell surface to 1 µm below the cell surface), the probability of gigaseal formation increases from 0 to 100% relatively linearly. When the pipette was pushed further into the cell, the gigaseal rate plateaus at 100% between 1 µm to 2.5 µm below the surface. As the pipette is advanced further, up to 7 µm below the cell surface, the gigaseal rate decays back to 0%. Each probability plotted in Figure 2b was calculated as the result of at least 4 attempts.

### 1.1.12 Synchronization of pipette-membrane distance

An example of the motion compensation process can be seen in Figure 3. Here, a 10 MΩ pipette is used to form a gigaseal with a thalamic cell after compensation for the cardiac and respiratory motion. The peak-to-peak cardiac current fluctuations were reduced by 89% (1.14 to 0.13 nA) and the peak-to-peak respiratory current fluctuations were reduced by 51% (2.35 to 1.16 nA). Following a suitable reduction in the current fluctuation amplitudes, a slight negative pressure (−30 mbar) was applied to form a gigaseal. The cell achieved a gigaseal in 3.7 s, and, importantly, achieved a high series resistance (400 MΩ) before the arrival of the next respiratory event. We surmise that a large series resistance, but not necessarily a gigaohm seal, is critical for mechanical stability during periods of high-amplitude motion.

**Figure 3:**
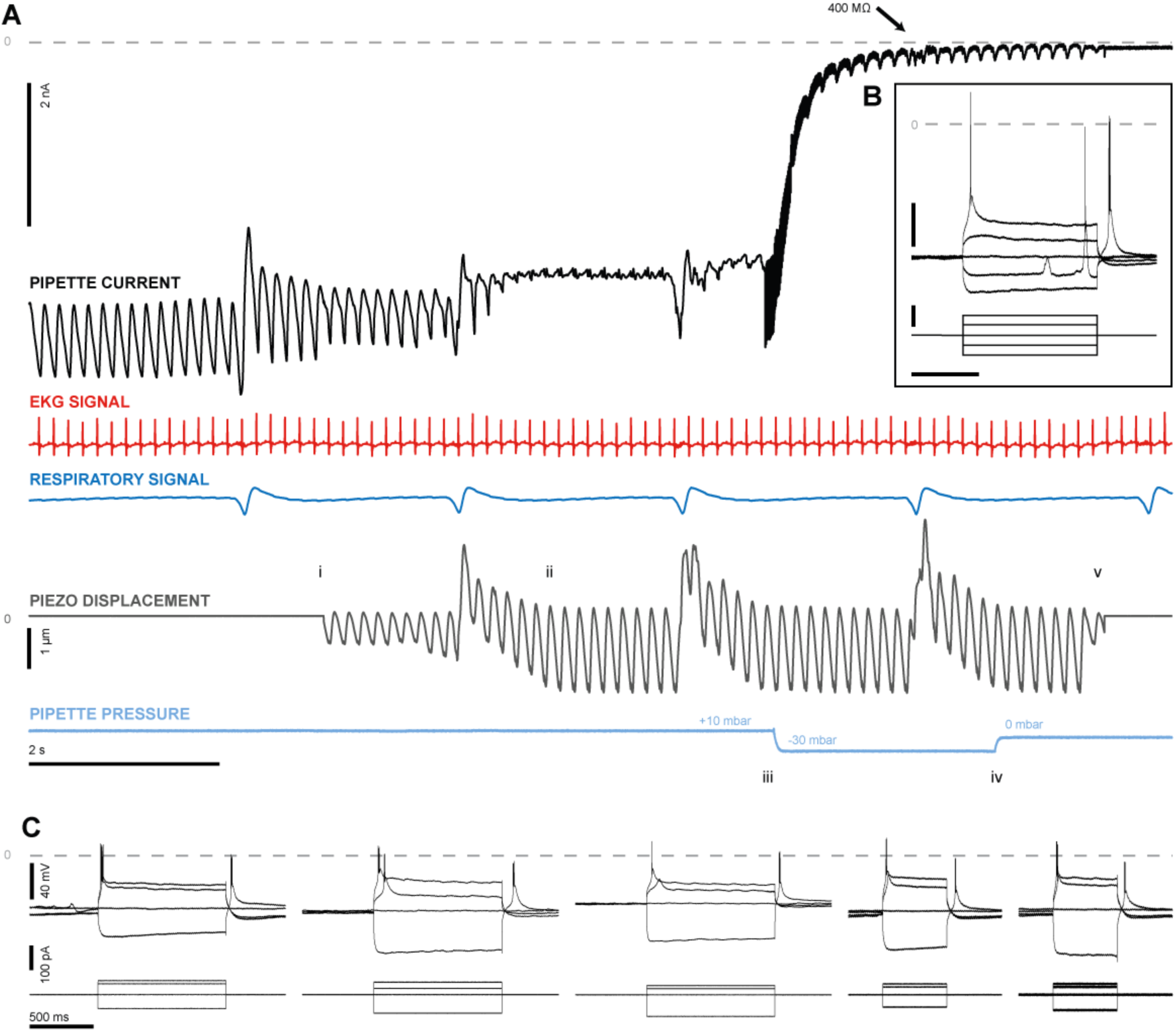
Motion compensation method applied to mouse thalamus *in vivo*. The recorded cell is held at -65 mV in voltage clamp mode and fluctuations in the measured current are recorded. Fluctuations in the measured current are indicative of a nearby cell moving with respect to the pipette tip. The EKG and respiratory sensor signals are used to determine physiological motion impulse responses. These impulse responses are used to drive the pipette axially using a high-resolution piezo stage, synchronize the motion of the pipette tip to the cell, and achieve a gigaseal as follows. i) The impulse responses are convolved with the live, thresholded EKG and respiratory input and used as a command signal for the piezo stage. ii) The impulse responses are manually scaled until the pipette current fluctuations are minimized, and the user sets the gigaseal mode to trigger after the next breath. iii) Suction is applied to the pipette and the gigaseal begins to form. The series resistance of the electrode is measured using Ohm’s law during gigasealing by applying a 128 Hz, 20 mV square wave. iv) The gigaseal has nearly formed and suction is removed. v) The gigaseal has formed and the seal is now mechanically stable. The motion compensation is halted. Following this procedure, break in can be performed automatically to perforate the membrane and achieve a whole-cell configuration. Inset shows current clamp recording from the same cell which demonstrates a burst spiking rebound following hyperpolarization, characteristic of thalamic neurons. Upper vertical scale bar, 20 mV. Lower vertical scale bar, 100 pA. Horizontal scale bar, 500 ms

To evaluate the success of the feedforward system at approximating the finite impulse response of physiological sources of motion in the brain, whole-cell patch clamp trials were performed in the cortex (n=7 mice, 70 trials) and in the thalamus (n=8 mice, 76 trials). Neuron hunting was performed as we published previously (Kodandaramaiah et al., 2012; Stoy et al., 2017). The forward progress of the pipette was halted when the pipette resistance increased above threshold and the peak-to-peak amplitude of the resistance fluctuations was above a user-determined threshold, indicating that the pipette was near a cell.

Once near a cell, motion compensation was attempted while measuring the pipette current in voltage clamp mode with a DC voltage applied (−65 mV). Unless otherwise stated, these results will discuss the reduction in the peak-to-peak current amplitude resulting from cardiac motion. In the thalamus, the peak-to-peak amplitude of the current fluctuations immediately prior to the motion compensation was 0.83 ± 0.56 nA and was reduced to 0.21 ± 0.18 nA (an average of 72.5% reduction in amplitude, when measured on a trial-by-trial basis). In the cortex, the peak-to-peak amplitude before motion compensation was 0.67 ± 0.41 nA and was reduced to 0.25 ± 0.13 nA (a trial-by-trial average of 57.5% reduction in amplitude). The initial peak-to-peak current amplitude was not significantly different between the thalamus and cortex (cortex: 0.67 ± 0.41 nA, thalamus: 0.83 ± 0.56 nA, p=0.326, Wilcoxon rank sum test). However, the average amplitude of motion compensation in the thalamus and cortex was 3.7 ± 1.42 µm and 1.7 ± 1.14 µm, respectively, and was found to be significantly different (p=3.62e-6, Wilcoxon rank sum test).Reducing the relative motion between the pipette and the cell in the thalamus resulted in a substantial increase gigaseal yield. Gigaseals were formed in 32/66 trials where regional pipette localization and neuron hunting were performed successfully. This is a significantly higher gigaseal yield in the thalamus achieved using only robotic navigation during regional pipette localization (RPL) (17/64 trials, See Figure 4, p=0.012, Fisher’s exact test, Stoy et al 2017), and is the highest reported *in vivo* gigaseal yield in the thalamus that we are aware of.

**Figure 4:**
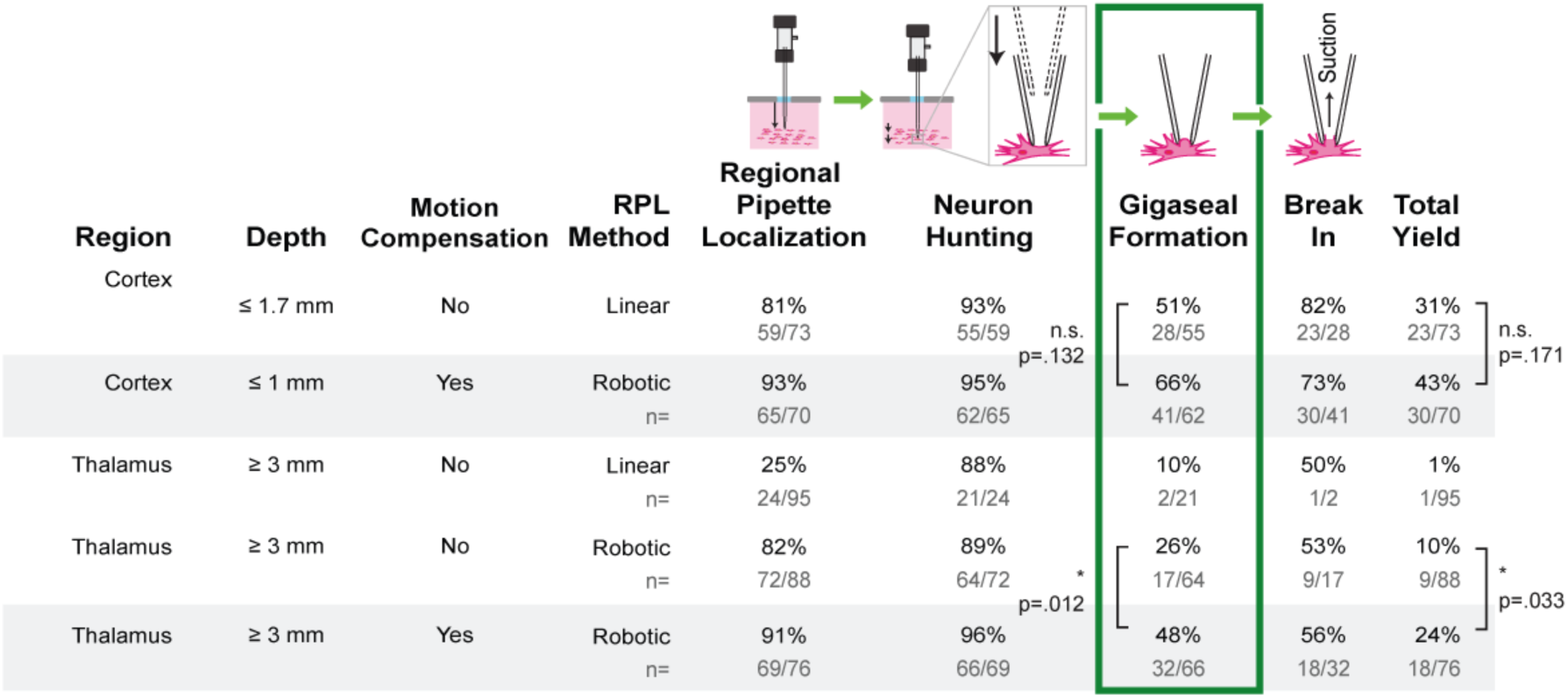
Results from automated whole-cell patch clamp experiments *in vivo*. Rows 1, 2: Cortex and Hippocampus. Results from (Kodandaramaiah et al., 2012) are compared with motion compensation trials in the cortex. Rows 3-5: Mouse thalamus whole-cell recording trials using linear (Kodandaramaiah et al. 2012, row 3) and robotic (Stoy et al. 2017, rows 4 and 5) regional pipette localization methods are compared with trials using motion compensation during the gigasealing stage. p values are calculated with Fisher’s Exact Test.

In the cortex, trials utilizing both motion compensation and robotic navigation achieved whole-cell configurations at a rate similar to Kodandaramaiah et al (See Figure 4, first row, p=0.171), and our gigaseal yield (66%, n=41/62) was similar to our previous efforts in the mouse cortex ((Kodandaramaiah et al., 2012): 51%, n=28/55, p=0.132, Fisher’s exact test).

Following the formation of a gigaseal, the break-in procedure is enacted, and the membrane is ruptured with one or more brief pulses of suction to form a whole-cell state. The whole-cell yield, for all pipettes inserted to a depth of 3+ mm in the thalamus using the methods described here, was 24% (n=18/76 attempts, Figure 4), which represents a significant improvement over our previous automated patching efforts in the thalamus (10%, n=9/88 attempts, p=0.033, Fisher’s Exact Test), and over all previously reported patching efforts in the thalamus to our knowledge.

### 1.1.13 Vibrissae pathway investigation with whisker and optogenetic stimulation

We demonstrated the efficacy of the motion compensation in an experiment on a preparation utilizing optogenetic and whisker stimulation and whole-cell patch clamp electrophysiology (Figure 5A). Sensory information is transduced at the whisker and sent through the trigeminal nucleus and thalamus until it reaches the primary sensory cortex. Each individual whisker has a discrete representation that is preserved along this trigeminal-thalamic-cortical pathway, making this system ideal for the study of sensory perception. In this experiment, the Nsmf-Cre line was crossed with the Ai32 mouse to express the channelrhodopsin, ChR2, in the primary whisker sensory thalamic region (VPM). By combining whole-cell recording and optical stimulation of the VPM axonal terminals, we could explore cortical circuitry while bypassing several sensory processing areas, including the trigeminal nucleus and the thalamus. This complex experiment required several days of preparation including intrinsic optical mapping and genotyping, and therefore required high yield cortical recordings. We found that our automated patch clamping system using motion compensation was able to efficiently record from the cortex.

**Figure 5:**
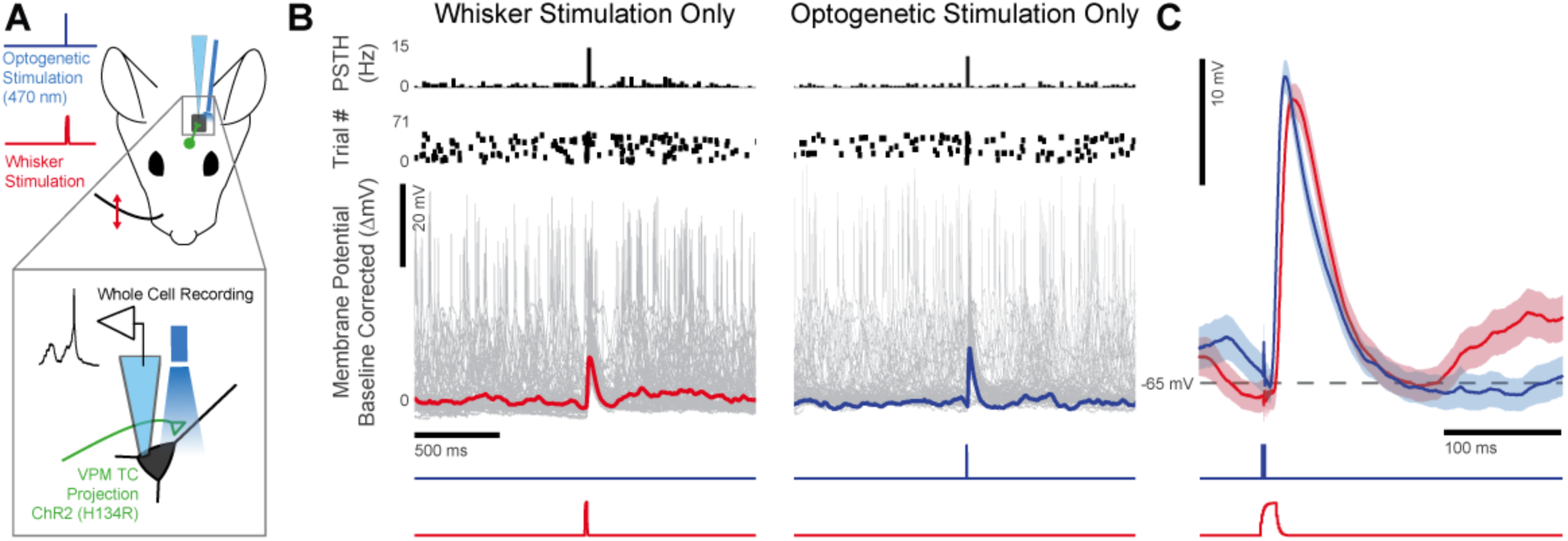
Results from cortical whole-cell patching with motion compensation. A) Schematic representation of the experimental setup. Whole-cell recording was performed in somatosensory cortex, layer 4 of an anesthetized Nsmf-CrexAi32 mouse. Whisker stimulation was performed by threading the C2 whisker into a capillary attached to a computer-controlled galvanometer. Light stimulus was delivered through a fiber-coupled 470 nm LED and stimulated axonal projections from the VPM with the fiber placed near the cortical surface. Trials of punctate whisker stimulus and light stimulus (1ms pulse) were embedded in background whisker stimulation (gaussian noise) or silence. B) Representative current-clamp recordings with the baselines subtracted have been stimulus-aligned. Gray traces indicate individual trials and red and blue traces indicate the median subthreshold activity (spikes have been removed and interpolated as described in Section Error! Reference source not found.). C) Detail of subthreshold responses to whisker stimulus (red) and light stimulus (blue). Shading indicates standard error of the mean (n=72 trials each).

We used these subthreshold recordings to look at timing differences of sensory stimuli and optogenetic stimulation. We observed a 3.9 ms delay between the onset of the mean subthreshold response on whisker stimulus trials compared to trials with optogenetic stimulation (Figure 5C). This delay was expected because optogenetic stimulation bypasses two synapses in the vibrissae pathway. Additionally, the firing pattern was similar between trials where VPM axon terminals were stimulated with light and trials where the whisker was stimulated, (Figure 5B). Similarly, when spikes were detected and removed by linear interpolation (1.5 ms before the spike peak, 4.5 ms following the spike peak), the subthreshold response dynamics were preserved. Specifically, the average full width half max value was similar (whisker: 36 ms, optogenetic: 30 ms) and the peak amplitude was also similar (whisker: 17 mV, optogenetic: 17.8 mV).

We performed 13 trials in 2 mice and achieved a whole-cell configuration in 8 attempts (61.5% success rate). In 7 trials, we recorded physiologically relevant signals for between 4-43 minutes (n=7; 4, 13, 14, 16, 35, 35, 43 minutes) and 5 of these cells were found to be whisker-responsive, and of those cells, 2 were also found to be responsive to optogenetic stimulation. An additional 2 whisker responsive cells were found to be responsive to optogenetic stimulation. Three of these recordings were terminated early following sufficient data collection so that more trials could be performed. This demonstrates that these recordings are not only stable, but that there is no damage to the cells caused by the addition of the motion compensation procedure.

## 1.3 Discussion

In this work, we have demonstrated 1) that there is a significant distance dependent relationship between the electrode tip and cell membrane which affects the likelihood of gigaseal formation *in vitro*, 2) that the distance between the membrane and the pipette fluctuates during the gigasealing step *in vivo*, 3) that by measurement of the pipette current and physiological signals, the impulse response can be rapidly estimated, 4) that by convolution of the impulse responses with incoming physiological signals, on-line compensation of the distance between the pipette and the cell membrane can be performed, and, critically 5) that this stabilization procedure results in a significantly higher gigaseal rate in deep subcortical nuclei, such as the thalamus. Importantly, this gigaseal rate is commensurate with previously reported yields for manual and Autopatching yields *in vivo* in the cortex (Kodandaramaiah et al., 2012). This method therefore opens the door to high-yield patch clamping studies throughout the brain at yields similar to those expected previously in only cortical layers (< 1 mm deep) *in vivo*.

The whole-cell recording yield is the product of the yield of the four stages of the patch algorithm (See Fig. 4). We note that there is still a low rate of successful break-in (in this work, 56% in the thalamus, 68% in the cortex), with deep patching that is irrespective of localization method, linear or robotic. We suspect that further optimizing the average distance between the pipette and the cell during the break-in stage may improve the success. We hypothesize that break-in dynamics (and gigasealing dynamics for that matter) are at least partially related to the tension in the membrane (Suchyna et al., 2009), and that there may be a similar optimal tension on a cell membrane for break-in. Additionally, we hypothesize that membranes may be more likely to rupture poorly when large-amplitude motion arrives during the application of a high-negative-pressure pulse. Further experiments can be performed investigate the relationship between respiratory event arrival times and the failure rate of break-in attempts.

In this study, we report significantly higher gigaseal yield when motion compensation is applied during whole-cell patching trials performed in the thalamus. When a cell moves with respect to a pipette, the distance between the pipette and the membrane contributes to the amplitude of the measured current. This relationship is known to be nonlinear (Rheinlaender & Schäffer, 2013). For example, when the average distance is greater than the pipette diameter, no change in current will be measured. When the distance between the cell and the pipette decreases, the flow of current to ground is impeded by the cell membrane. As the cell nears the pipette during its modulated position cycle, we expect the flow of current to decrease, and as it moves further than the pipette diameter away, we expect the current to be rectified to the original pipette current. Finally, if the pipette has pressed into the cell far enough, we would again expect the pipette current to be rectified, indicating that the membrane is being pulled against the pipette (the cell is nearly impaled in this configuration). We did not notice that there was current rectification immediately prior to gigasealing on any of the trials performed in this study. Rather, the pipette current was slightly reduced (approximately 5-10% of the original current). This suggests that the pipettes were slightly indenting the surface of the cell’s membranes in this study, immediately prior to gigasealing. However, this is the same decrease in current that we have used in all previous studies with the Autopatcher, both in the cortex and the thalamus (Kodandaramaiah et al., 2012; Stoy et al., 2017). Further studies may be performed to investigate the success rate due to average distance from the pipette tip to the membrane *in vivo* and *in vitro* in brain slices. Specifically, a study of distance dependent gigasealing rate can be performed in neurons in slices of brain tissue to more closely reflect the deformability of the cells and their surroundings, as opposed to the cells plated on stiff substrates used in this study. We still expect that there will be an ‘inverted U’ shaped relationship of gigaseal probability with respect to pipette-membrane distance, similar to Figure 2. Regardless of preparation, pipettes at large distances from cell membranes will never form gigaseals and pipettes that are pushed too far into the cells will damage the membrane. Therefore, we expect that the region of high yield gigaseal formation will be larger, but not indefinitely. Further, the long-term health of cells that have been greatly indented during gigaseal formation is of concern. Any study that seeks to establish a robust distance dependent gigaseal rate for cells in brain slices should take this factor into account and measure the cell’s resting membrane potential and the longevity of the whole-cell recording.

The motion compensation hardware and software described here is part of an automated suite of tools for neuroscientists that enables the recording of whole-cell electrophysiology in previously difficult-to-access tissue (Stoy et al., 2017). Additionally, the combination of this motion compensation procedure and robotic localization during regional pipette localization with pipette cleaning and reuse (I Kolb et al., 2016) enabled full automation of demanding experiments in highly valuable tissue. Together, this constellation of tools represents empowering technology for the field of neuroscience to be able to gather repeatable, high-throughput, high-quality data throughout the brain from fewer animals with minimal human intervention.

## CREDIT STATEMENT

**William Stoy:** Conceptualization, Methodology, Software, Investigation, Formal Analysis, Writing-Original draft preparation, Writing - Review & Editing **Bo Yang:** Methodology, Writing-Reviewing and Editing **Ali Kight:** Methodology, Writing-Reviewing and Editing **Peter Borden:** Conceptualization, Methodology, Resources, Writing- Reviewing and Editing **Nathaniel Wright:** Conceptualization, Methodology, Writing- Reviewing and Editing **Garrett Stanley:** Supervision, Writing- Reviewing and Editing **Craig Forest:** Conceptualization, Supervision, Writing- Reviewing and Editing

## SUPPLEMENTARY INFORMATION

### S1.1 Optimization of *impulseest*

Parameters for *impulseest* were tuned to minimize the time to calculate an impulse response and maximize the accuracy of the estimation. This process was performed by selecting segments of pre-recorded data (training data), sampling at 200 Hz, in integer lengths from 4-15 seconds with random offsets and selecting various model orders (64, 128, 256). The cardiac and respiratory impulse responses were calculated 10 times for each combination of parameters. The total time to produce the impulse response estimations was recorded (Total time = recording time + impulse estimation time). Additionally, the quality of fit of the model was assessed on a different set of training data. Specifically, the resulting impulse responses were convolved with the thresholded cardiac and respiratory signals and the sum of squared errors between the convolved signal and the true current trace was calculated.

The results of this optimization can be seen in Supplementary Figure 2. We chose a model order of 128 instead of 64 because the marginal time required to calculate the impulse response function was negligible and because at a sampling rate of 200 Hz, the impulse response length is 0.65 seconds, which we determined was sufficient to capture all respiratory dynamics (data not shown). Additionally, we chose a training data length of 8 seconds because marginal increases in training data length beyond this value did not confer significantly better fit (lower SSE from Supplementary Figure 2). Using these parameters, the FIR model was computed using the training data in an average of 0.29 ± 0.02 seconds.

Although the work of Michale Fee in his 2000 paper entitled, “Active Stabilization of Electrodes for Intracellular Recording in Awake Behaving Animals”, provides an important conceptual basis for the above method, there are a number of key differences present in the work shown here (Summarized in Supplementary Table 1). Fee computed the impulse response using the Fourier transform and standard spectral analysis techniques with (approximately) the inverse Fourier transform of the transfer function. The main advantage of Fee’s method is that, computationally, it is exceptionally fast (1 ms in our implementation). In contrast, we used a least-squares estimation, implemented with MATLAB’s *impulseest* function. The *impulseest* function proceeds with a recursive least-squares estimation, producing single-shot impulse responses that, when convolved with the physiological input signals, has high correlation to the measured system, as evidenced by the 71% average reduction in current fluctuation amplitude. The computation time of this recursive model is significantly longer (∼ 0.5 s) than the spectral analysis techniques used by Fee. However, the result of this single-shot impulse estimation is that it takes an overall shorter time to apply the impulse response than Fee due to the reduced recording time. Fee used an iterative process to update the cardiac and respiratory impulse responses. Each computation requires 2-4 seconds and is updated at least once, sometimes up to 3 times. This results in a total recording / computation time of 8-24 seconds. Reducing the time to apply the motion compensation is likely critical to the success of gigaseal formation and to the health of the cell following break-in. When the pipette is in proximity to a cell before gigaseal formation is attempted, it is under positive pressure. This pressure causes flow of intracellular fluid out of the pipette tip and alters the ionic composition of the extracellular space. Prolonged perfusion of this space with intracellular solution can damage cells, precluding successful gigaseal formation or reducing the longevity or physiological relevance of whole-cells that are achieved. Additionally, expulsion of unnecessary intracellular solution will degrade the image quality with high background fluorescence or off-target staining if the pipette contains a fluorophore or biocytin for single cell labelling.

### S1.2 Transgenic mice

We used transgenic mice to express the light sensitive protein channelrhodopsin (ChR2) into the lemniscal thalamic projecting neurons using cre/cre-dependent knock-in breeding. Specifically, we used the Tg(Gsat-NR133-Nsmf-Cre) mouse line to express Cre in the primary somatosensory thalamic regions of the VPm/VPL, and crossbred this mouse line with the Ai32(RCL- ChR2(H134R)/EYFP) mouse line (JAX Laboratories, Stock No#:024109). Mice were genotyped Ttransnetyx, Inc) to confirm the presence of the NR133-Nsmf, YFP, and Rosa genes.

### S1.3 Barrel cortex mapping with intrinsic imaging

The mouse primary somatosensory cortex is responsive to physical stimuli and touch sensations on the mouse’s body. A large area of representation in this region is the barrel cortex, so called for its discrete, high density cell clusters approximately 100 µm in diameter called barrels, each representing a single whisker.

Prior to each experiment, the location of the C1 and C2 whisker barrel was determined using Intrinsic Signal Optical Imaging (ISOI). Each mouse’s barrel cortex was imaged through either intact or thinned skull using a wide-field imaging system to measure cortical spatial activity (184×123 pixel CCD Camera, MiCam02HR SICMedia, Ltd). During all imaging experiments, isoflurane anesthesia levels were lowered to approximately 1%. The cortex was imaged at 10 Hz with a field of view of 4×3mm with a total of a 1.6 Magnification (48 pixels/mm). We used a green (535nm LED) excitation light projected onto the cortical surface that has a high overlap with the hemodynamic absorption spectrum. Collected light was filtered with a set of dichroic mirrors (Bandpass 475/150 nm and Longpass 495 nm, Semrock,Inc) and a bandpass emission filter between wavelengths of 520/35 nm (Semrock, Inc). The imaging system was focused at approximately 300 µm below the cortical surface to target cortical layer 2/3. In order to evoke a cortical intrinsic response, the whisker was repetitively stimulated at 10Hz for 6 seconds.

The activity map and co-registered blood vessel map generated during the ISOI session is shifted, rotated and deformed to match it with the vasculature observed on the experiment day. The activity map indicates the region of activity for the C1 whisker and is targeted by performing a craniotomy over the location of the dark spot seen in the center of the image.

Imaging of a single barrel is performed on a plane parallel to the brain surface; however, the pipette axis is 12.5° off of this axis. To compensate for this angular difference, the pipette is brought to the surface of the brain at the center of the active region, identified by aligning the co-registered ISOI map to the vasculature (See Supplementary Figure 3), and then moved approximately 89 µm medial (See Supplementary Figure 4).

### S1.4 Optical stimulation

Axonal projections terminating in the barrel cortex were stimulated with a blue light (470 nm, M470F3, THORLABS), which was driven at 1.2 Amps with a THORLABS LEDD1B driver. The LED was coupled to a 400 µm diameter multimodal optic fiber which was positioned using a manual micromanipulator to within 200 µm of the pial surface above the craniotomy. The LED was triggered in TTL mode using a command signal from the cDAQ-9174.

### S1.5 Whisker stimulation

Whisker stimulation was performed using a galvo (galvanometer optical scanner model 6210H, Cambridge Technology) with a custom milled aluminum fixture with holes to permit the reliable stimulation of a single whisker. The stimulator was placed on the right side of the mouse’s face 5 mm from the whisker pad. The rotating arm of the galvo-motor was arranged such that the mean whisker position during noise stimulation was its resting point and the stimulus was delivered along the rostro-caudal axis. The galvo was controlled using an analog output signal from the cDAQ-9174 and the galvo position was recorded on the Digidata 1550 Digitizer. Whisker punctate stimuli were delivered at 1200 µm. White noise was delivered to the whisker on half of the trials with punctate whisker stimulation and half of the trials with optogenetic stimulation.

**Supplementary Figure 1:**
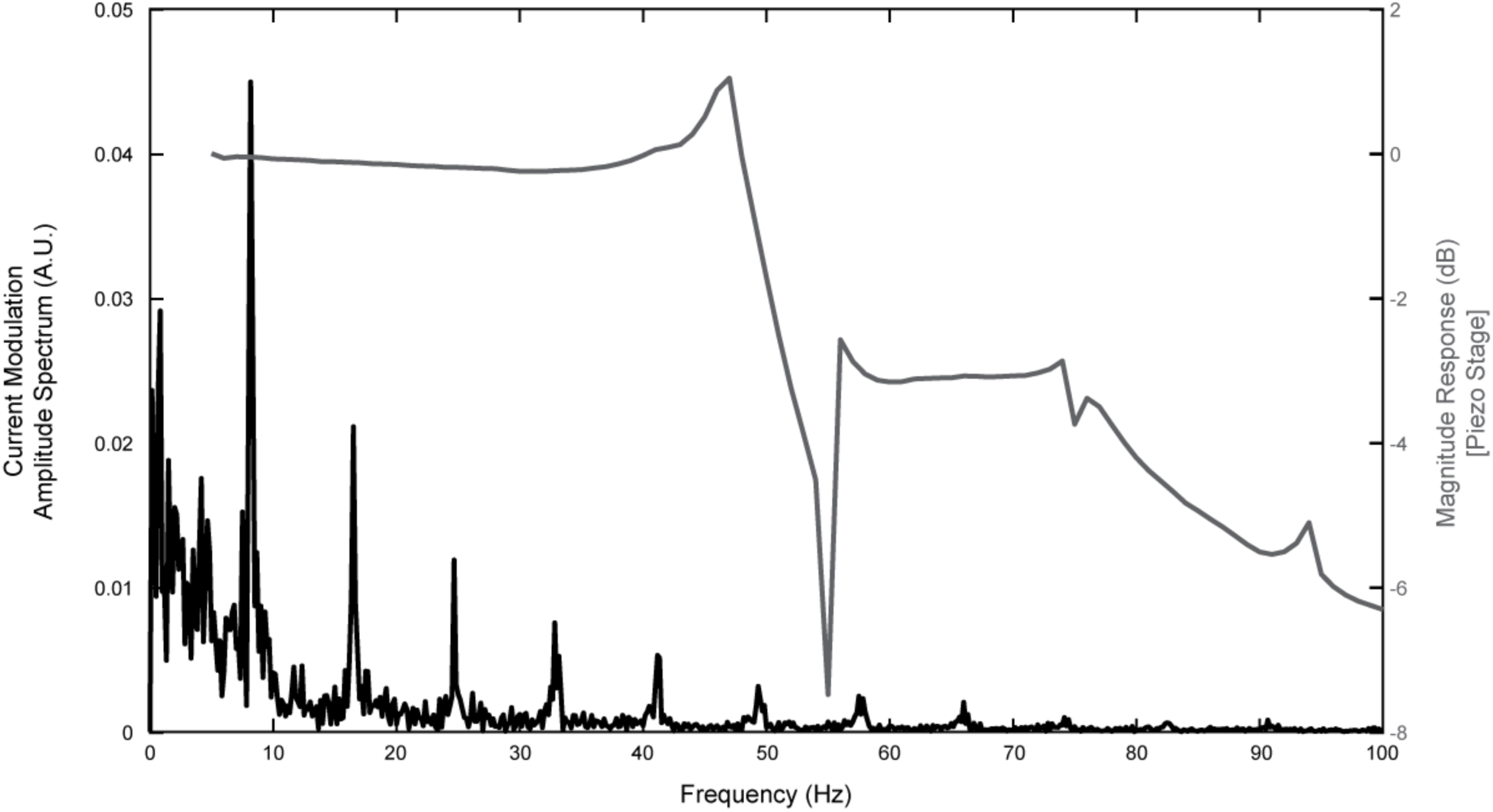
Spectral analysis of current modulation and the piezo stage. The amplitude spectrum of a recording of 10 seconds of current modulation by respiratory and cardiac motion (blue) is shown along with the magnitude response of the piezo stage used in this study to a sinusoidal frequency sweep at 1 V_pp_ (corresponding to 10 µm peak-to-peak). The resonant peak of the piezo stage is at 47 Hz.

**Supplementary Figure 2:**
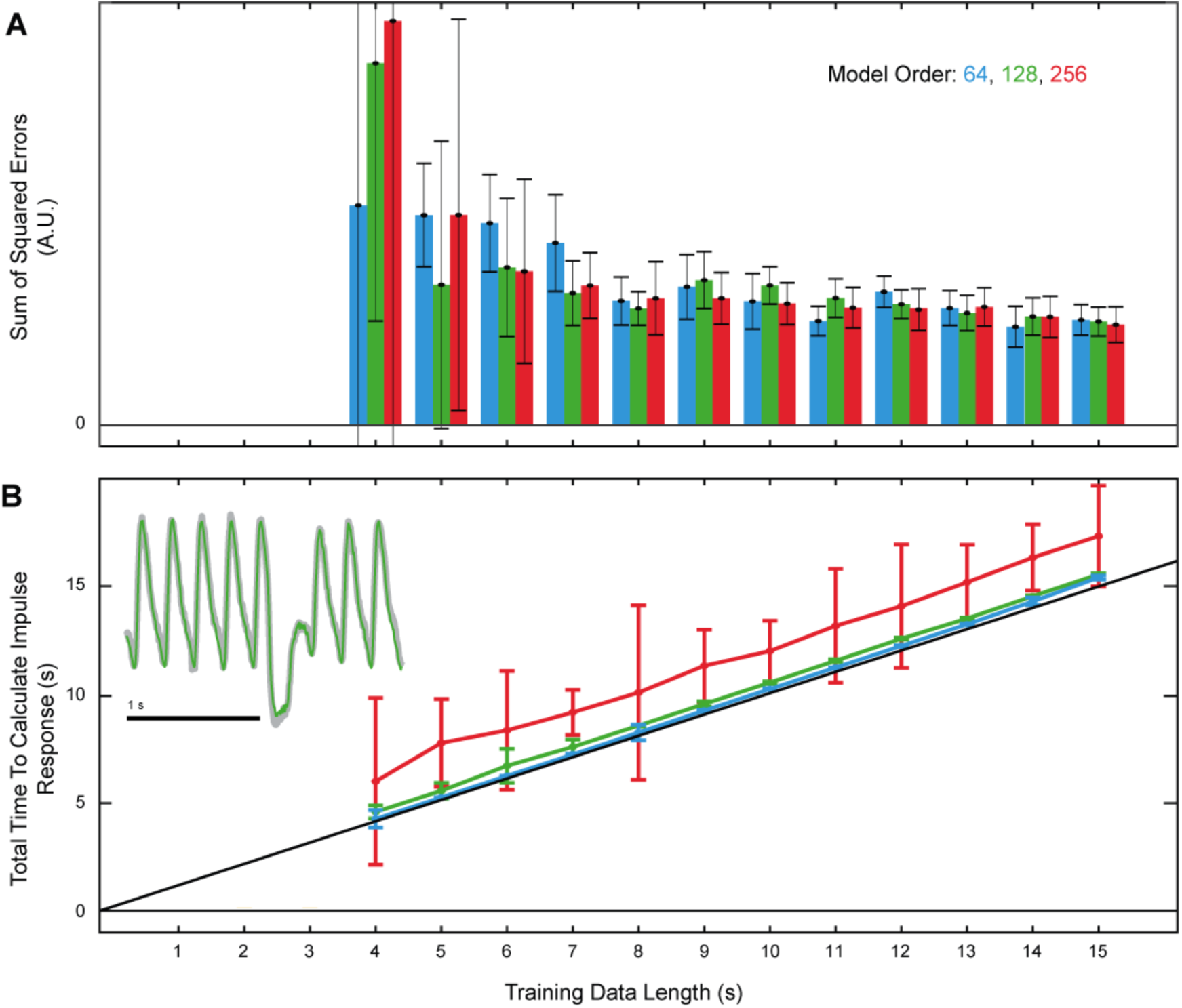
Optimization of *impulseest* parameters. Various training data lengths and model orders were selected. Each combination of parameters was run 10 times. A) The quality of the fit (SSE, lower is better), and B) total time to calculate impulse response are plotted. Total time to calculate impulse response is the sum of training data length and the impulse response calculation time. Diagonal line in lower graph is the unity line. Inset: example of cardiac and respiratory filters (order: 128, training data length: 8 s) convolved with corresponding pulse trains. The result of the convolution is plotted (green) over corresponding current trace (gray).

**Supplementary Figure 3:**
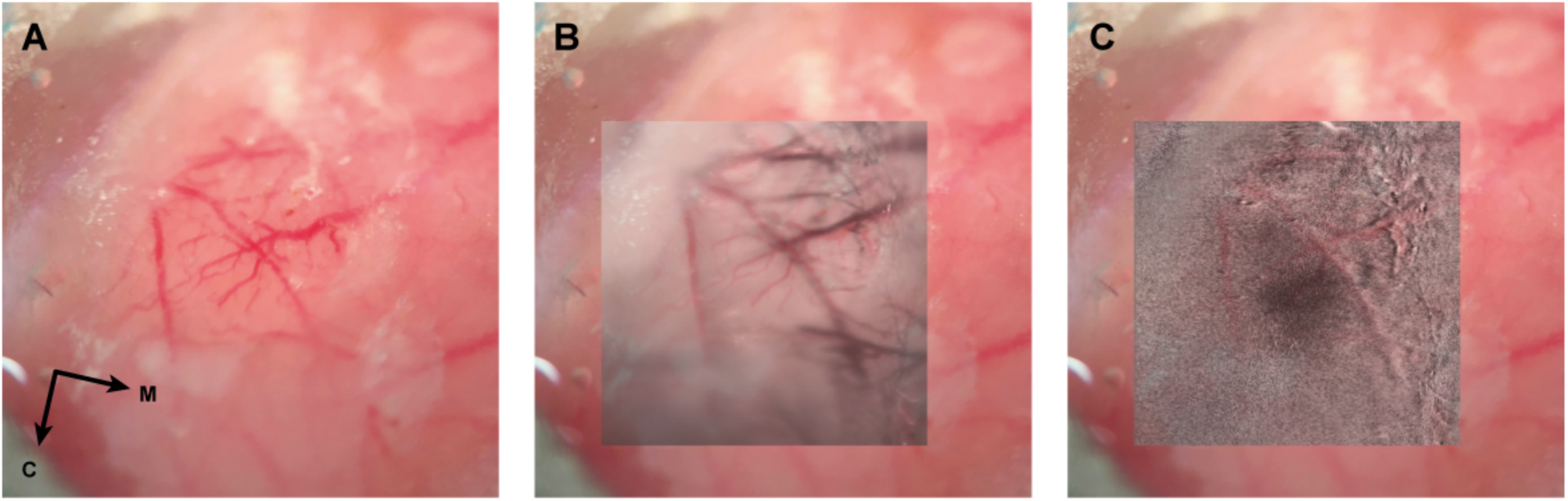
Alignment of intrinsic signal optical imaging (ISOI) signal with vasculature. Intrinsic images were captured 3 days prior to electrophysiology experiment. Here, the C1 whisker was rapidly stimulated and images were captured under red light (650 nm) before and after the stimulation and averaged over several trials. Darker regions indicate local consumption of blood oxygen, a proxy of activity. A co-registered blood vessel image is taken under 550 nm light. A) Cortical vasculature can be seen through the thinned skull preparation under perfusion of saline. B) Blood vessel map is shifted, rotated and deformed to match it with the vasculature observed on the experiment day. C) The co-registered activity map indicates the region of activity for the C1 whisker and can easily be targeted by performing a craniotomy over the location of the dark spot seen in the center of the image.

**Supplementary Figure 4:**
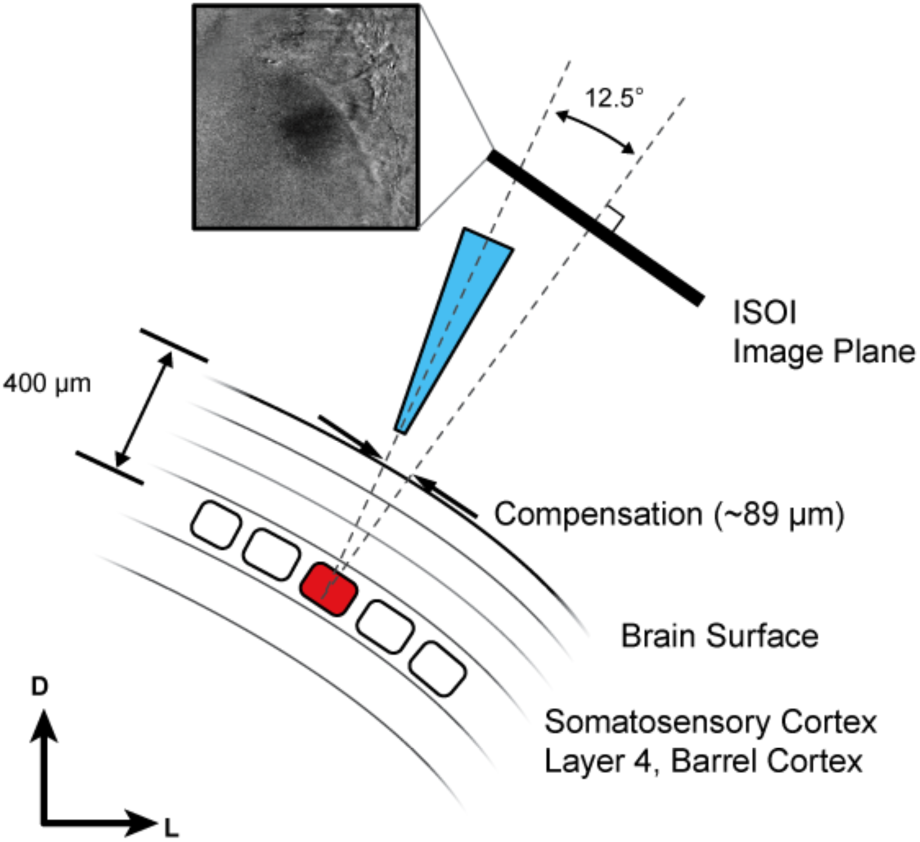
Compensation of angular difference between pipette axis and ISOI normal. The pipette axis is 12.5° from the axis normal to the ISOI image plane. The location of the barrel of interest is determined with respect to the pipette axis by trigonometry. The pipette (shown in blue here) is then brought to the surface of the brain in the center of the area of activity determined by the ISOI session and then moved approximately 89 µm medial to ensure that the pipette passes through the barrel of interest (red rectangle).

**Supplementary Table 1:** Comparison of electrode-based motion compensation methods. Note: total computation time in Michale Fee’s work was determined by extrapolating from results in the text. The cardiac FIR filter is computed and then the respiratory FIR filter is computed. Each computation requires 2-4 seconds and is iteratively updated at least once, sometimes up to 3 times. This results in a total recording / computation time of 8-24 seconds.

|  | Impulse Response<br>Computation Method | Total Recording /<br>Computation Time | IRF scalar z/i<br>Calculation | Motion<br>Reduction |
| --- | --- | --- | --- | --- |
| Michale Fee<br>(2000) | Spectral Analysis<br>$h = F^{-1} \left( \frac{F(y)}{F(x)} \right)$ | 8 – 24 s recording<br>$\ll 0.1$ s<br>computation | 80 Hz sine<br>wave (1 s) | ‘80% or<br>more’ |
| This work | Least-Squares Estimation<br>$y = Xh + e$<br>$h = yX^{-1}$ | 8 s recording<br>$0.55 \pm 0.36$ s<br>computation | Manual<br>(1 s) | $71 \pm 13\%$<br>reduction |

